# Oncogenic Activation of AKT by MAPK6

**DOI:** 10.1101/2020.09.23.309518

**Authors:** Qinbo Cai, Wei Wang, Bingning Dong, Wolong Zhou, Tao Shen, David D. Moore, Chad J. Creighton, Feng Yang

**Affiliations:** Department of Molecular and Cellular Biology, Baylor College of Medicine, Houston, Texas 77030 USA; Department of Medicine, Baylor College of Medicine, Houston, Texas 77070 USA; Dan L Duncan Comprehensive Cancer Center, Baylor College of Medicine, Houston, Texas 77030 USA

## Abstract

Mitogen-activated protein kinase 6 (MAPK6) is an atypical MAPK closely related to MAPK4. We recently reported that MAPK4 can promote cancer by activating the Protein Kinase B (PKB/AKT) pathway of cell growth and survival. Here we report that MAPK6 overexpression also activates AKT to induce oncogenic outcomes, including transforming “normal” human epithelial cells into anchorage-independent growth and enhancing cancer cell growth. Knockdown of MAPK6 inhibited cancer cell growth and xenograft growth, supporting the tumor-promoting activities of endogenous MAPK6. Unlike MAPK4, which binds AKT through its kinase domain and phosphorylates AKT at T308, MAPK6 interacts with AKT through its C34 region and the unique C-terminal tail and phosphorylates AKT at S473 independent of mTORC2, the major AKT S473 kinase. MAPK6 overexpression is associated with decreased overall survival and the survival of lung adenocarcinoma, mesothelioma, uveal melanoma, and breast cancer patients. We conclude that MAPK6 can promote cancer by activating AKT and that targeting MAPK6 may be effective in human cancers.

## Introduction

MAPK6 is considered an atypical MAPK because its activation loop contains a Ser-Glu-Gly (SEG) motif compared to the conserved Thr-X-Tyr (TXY) motif in the canonical MAP kinases (1). In the canonical pathway, MAPK activation is dependent on the MAPK kinase kinase (MAPKKK or MAP3K) and MAPK kinase (MAPKK or MAP2K) signaling cascade (2, 3). In contrast, MAPK6 appears to be phosphorylated at S189 of the SEG motif and activated by the group I p21-activated protein kinases (PAK1/2/3) (4, 5). MAPK6 can phosphorylate several protein substrates, including MAP kinase-activated protein kinase 5 (MAPKAPK5 or MK5) at T182 (6, 7), steroid receptor coactivator 3 (SRC-3) at S857 (8), and tyrosyl DNA phosphodiesterase 2 (TDP2) at S60 (9).

While the roles of canonical MAPKs, such as MAPK3/1 (ERK1/2), MAPK11-14 (p38β/γ/δ/α), and MAPK8-10 (JNK1/2/3), are well established in carcinogenesis and tumor progression (2, 3), the role of MAPK6 in human cancers remains unclear. The best described tumor promoting activity of MAPK6 is enhancing the migration/invasion/metastasis of cancer cells, including those of lung, head and neck, breast, and cervix, as well as the human umbilical vein endothelial cells (HUVECs) and vascular smooth muscle cells (VSMCs) (8, 10-15). MAPK6 also promoted HUVEC, VSMC, and cervical cancer cell growth (10, 13, 14). In contrast, MAPK6 was reported to repress fibroblast cell cycle progression (16), melanoma and intrahepatic cholangiocarcinoma cell proliferation, and melanoma cell migration (17, 18), all of which support a tumor suppressor function. Thus, MAPK6 function in human cancers, especially its roles in regulating cancer growth, remains elusive.

MAPK4 is another atypical MAPK most closely related to MAPK6. Human MAPK4 and MAPK6 both contain an N-terminal kinase domain of 73% identity, a C34 region of nearly 50% identity, and a C-terminal extension that is not conserved (1). We recently reported that MAPK4 promotes cancer by functioning as a non-canonical AKT kinase independent of PI3K (19). This raised the question of whether AKT is also a substrate of MAPK6 and whether MAPK6 possesses tumor-promoting activity by activating AKT.

In this study, we report that MAPK6 overexpression induces oncogenic outcomes, including transforming “normal” prostate and mammary epithelial cells into anchorage-independent growth and enhancing cancer cell growth in an AKT-dependent manner. MAPK6 directly binds to AKT and phosphorylates AKT at S473. Unlike MAPK4 binding AKT through a CD motif in the kinase domain, MAPK6 interacts with AKT through the C34 region and the unique long C-terminal tail. We conclude that MAPK6 can promote cancer by activating AKT and that targeting MAPK6 may provide a novel therapeutic approach for cancer.

## Results

### MAPK6 overexpression transforms epithelial cells into anchorage-independent growth and promotes tumor cell growth

Previous studies have reported both tumor-promoting and tumor-inhibiting activities of MAPK6. To assess MAPK6 function, we first overexpressed it in the “normal” human prostate epithelial PNT1A cell line and breast epithelial MCF10A cells in a Dox-inducible manner (Figure 1A). We then assessed MAPK6 ability to transform these cells into anchorage-independent growth in the soft-agar assays. Similar to the robust activities of the closely related MAPK4 in transforming PNT1A cells (19), MAPK6 overexpression transformed both PNT1A cells and MCF10A cells into anchorage-independent growth (Figure 1B). In accord with an oncogenic activity of MAPK6, MAPK6 overexpression also potently promoted the growth, including the clonogenic growth of PNT1A and MCF10A cells (Figure 1, C and D).

**Figure 1.**
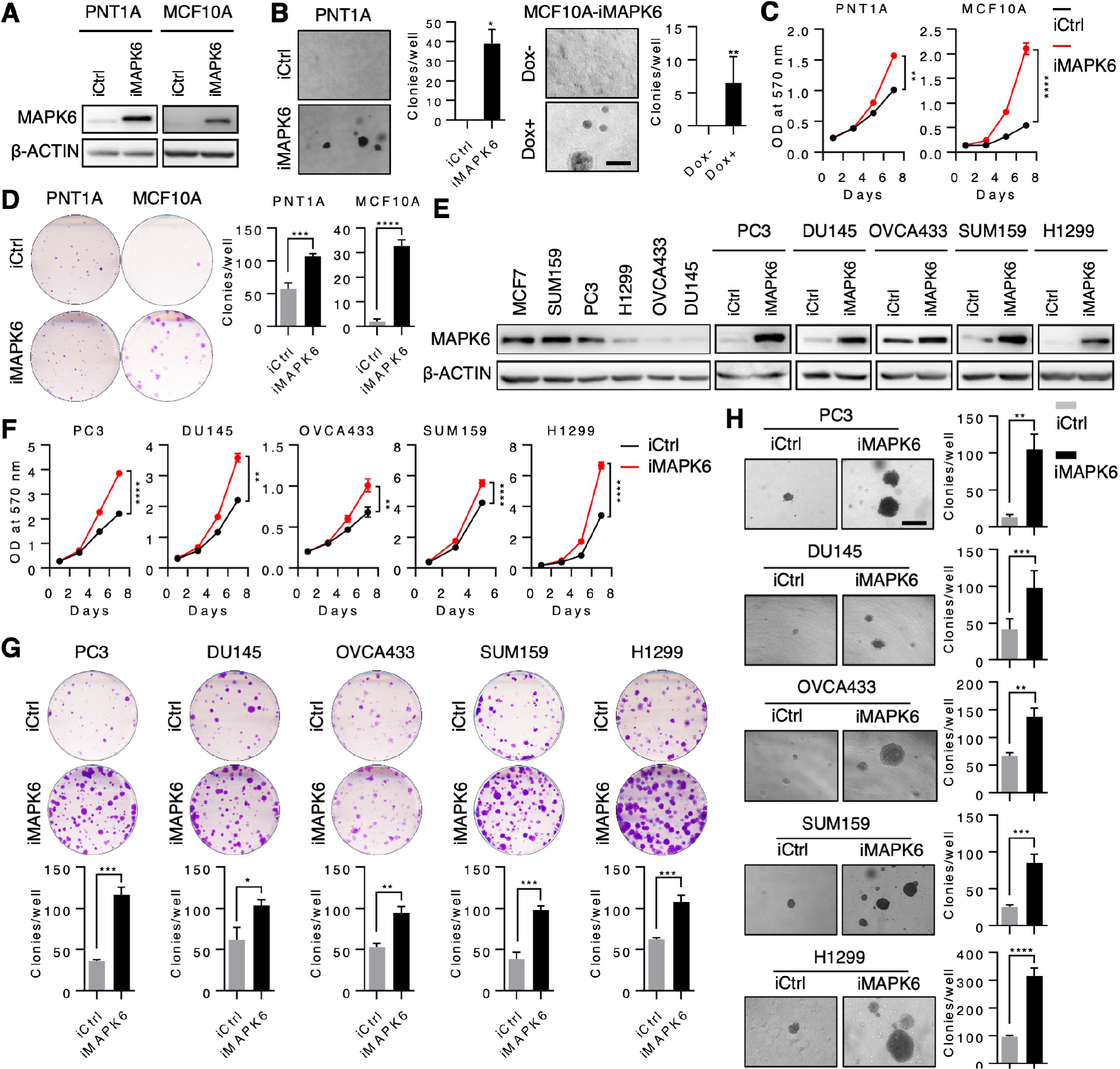
MAPK6 overexpression promotes tumor cell growth. (**A**) Western blots on doxycycline (Dox)-induced overexpression of MAPK6 in the engineered PNT1A and MCF10A cell lines. iMAPK6: Dox-inducible expression of MAPK6. iCtrl: control. (**B**) Soft-agar assays on MAPK6 transforming the engineered PNT1A and MCF10A cells into anchorage-independent growth. Shown are PNT1A-iMAPK6 and PNT1A-iCtrl cells induced with 1 µg/ml of Dox (left panels) and the MCF10A-iMAPK6 cells treated with (+) or without (-) Dox (right panels). Label: 500 μm. (**C**) Proliferation assays on the engineered PNT1A and MCF10A cells with Dox-induced overexpression of MAPK6 (iMAPK6) or control (iCtrl). Quantification data are shown as mean ± SD with indicated *P* values calculated by Two-way ANOVA. (**D**) Plate colony formation assays on the cell lines as described in (C). (**E**) Western blots on the endogenous expression and Dox-inducible overexpression of MAPK6 in different cancer cell lines. (**F**) Proliferation assays on the engineered PC3, DU145, OVCA433, SUM159, and H1299 cells with Dox-induced overexpression of MAPK6 (iMAPK6) or control (iCtrl). Quantification data are shown as mean ± SD with indicated *P* values calculated by Two-way ANOVA. (**G**) Plate colony formation assays on the cell lines as described in (F). (**H**) Soft-agar assays on the cell lines as described in (F) and (G). For inducible overexpression of MAPK6, 1 μg/ml Dox was used in all the above Dox-associated experiments. Label: 500 μm. Quantification data are shown as mean ± SD with indicated *P* values calculated by Student’s t-test except as otherwise indicated. iCtrl: Dox-inducible control. iMAPK6: Dox-inducible overexpression of MAPK6. *: *P* < 0.05. **: *P* < 0.01. ***: *P* < 0.001. ****: *P* < 0.0001. Data are representative of at least 3 independent experiments.

MAPK6 has been reported to exhibit tumor-promoting activities by enhancing tumor cell migration/invasion, but not growth (8). In sharp contrast, our data (Figure 1 A-D) support a growth-promoting role of MAPK6 in cancer cells. To further assess MAPK6 function in human cancer cells, we next examined the impact of Dox-inducible MAPK6 overexpression in several diverse human cancer cell lines with low to high endogenous levels of MAPK6, including the prostate cancer PC3 and DU145, ovarian cancer OVCA433, breast cancer SUM159, and non-small cell lung cancer H1299 cells (Figure 1E). Dox-induced MAPK6 expression significantly promoted the growth, including the clonogenic growth and the anchorage-independent growth of all these tumor cells (Figure 1, F-H).

To further investigate the biological function of endogenous MAPK6, we performed siRNA and/or Dox-induced shRNA knockdown of MAPK6 in the MAPK6-high or MAPK6-medium MCF7, SUM159, PC3, and H1299 cells (Figure 2A). In agreement with the gain-of-function studies, shRNA or siRNA-induced knockdown of MAPK6 significantly inhibited cell growth in the proliferation and clonogenic assays (Figure 2, B and C; Supplementary Figure 1, A and B). Dox-induced shRNA knockdown of MAPK6 also greatly inhibited the anchorage-independent growth of all these cells *in vitro* and H1299 xenografts growth *in vivo* (Figure 2, D and E). Altogether, our data support a tumor growth promoting function of MAPK6.

**Figure 2.**
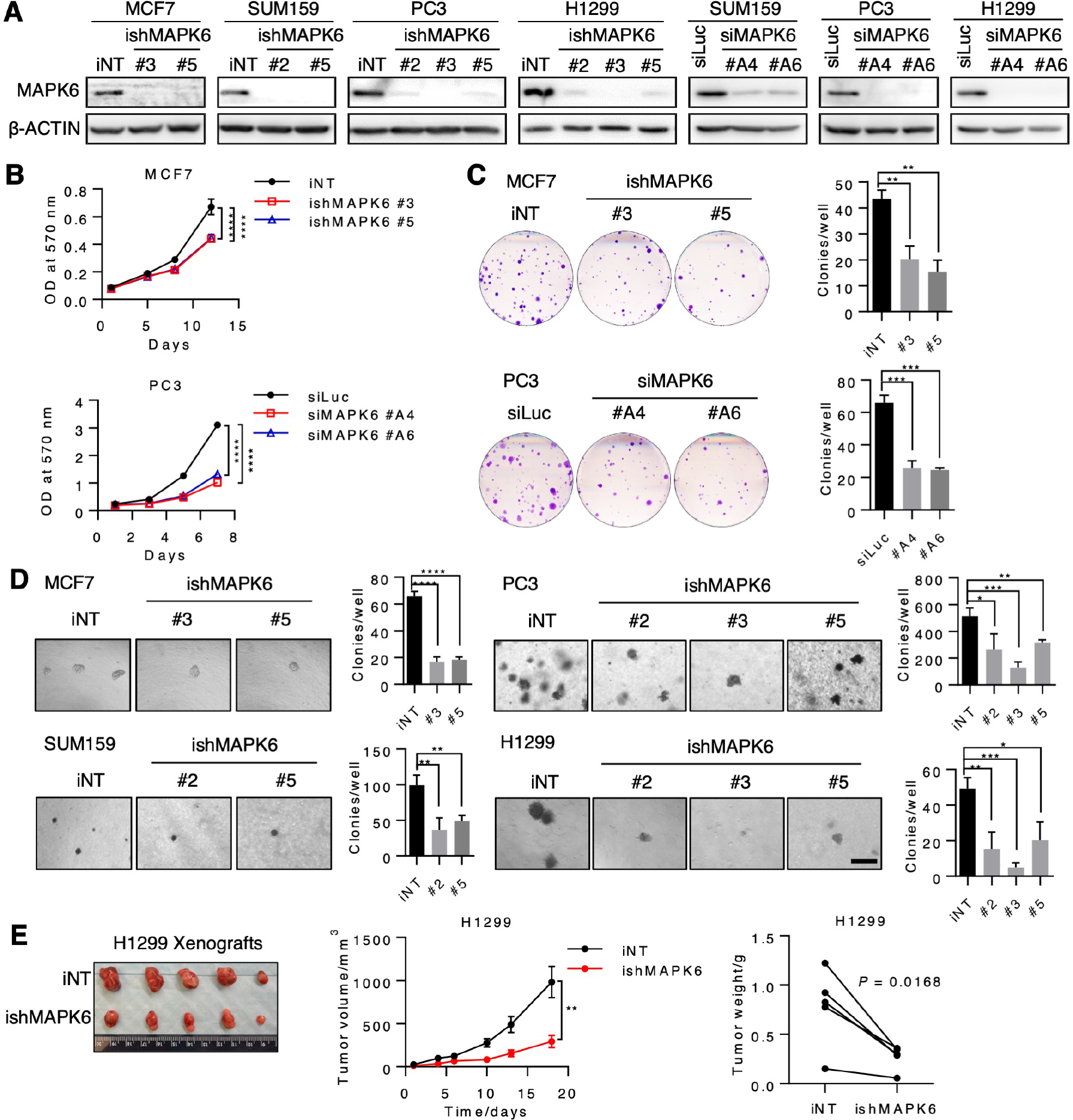
Knockdown of MAPK6 inhibits tumor growth. (**A**) Western blots on Dox-induced knockdown of MAPK6 by shRNA (ishMAPK6) or transient knockdown of MAPK6 by siRNA (siMAPK6) in cancer cell lines. iNT: inducible nontargeting control. siLuc: siRNA against luciferase as control. (**B**) Upper Panel: Proliferation assays on the Dox-induced engineered MCF7 cells with Dox-inducible knockdown of MAPK6 (ishMAPK6 #3 and #5 and iNT control). Lower Panel: Proliferation assays on the PC3 cells 48 hours after being transfected with siRNAs against MAPK6 (siMAPK6 #A4 and #A6) or control (siLuc). Quantification data are shown as mean ± SD with indicated *P* values calculated by Two-way ANOVA. (**C**) Plate colony formation assays on the cells as described in (B). Data shown are mean ± SD, Student’s t-test. (**D**) Soft-agar assays on the cells as described in (B) and (C). Data shown are mean ± SD, Student’s t-test. 4 μg/ml Dox was used for the Dox-inducible knockdown of MAPK6 for all the Dox-associated *in vitro* experiments. Label: 500 μm. Data are representative of at least 3 independent experiments. (**E**) Dox-inducible knockdown of MAPK6 significantly represses H1299 xenograft growth in SCID mice. A quantity of 1 × 10^6^ iNT or ishMAPK6 cells were s.c. injected into the lateral flanks of SCID mice (iNT: left side; ishMAPK6: right side). Mice began receiving 4 mg/ml Dox and 5% sucrose in drinking water on the day of tumor inoculation. Tumors were harvested as indicated. Left panel: image of tumors. Middle panel: growth curve of tumors (mean ± SEM, Two-way ANOVA test). **: *P* < 0.01. Right panel: tumor weight (paired Student’s t-test). Paired xenografts represent an iNT xenograft (on the left lateral flank) vs. an ishMAPK6 xenograft (on the right) within the same mouse. *P* = 0.0168. Data are representative of two independent experiments.

### MAPK6 activates AKT

MAPK6 is closely related to MAPK4. Since MAPK4 can directly activate AKT to promote tumor growth (19), we examined whether MAPK6 also activates AKT. To directly compare MAPK4 and MAPK6 activities in regulating AKT phosphorylation, we co-transfected FLAG-tagged AKT together with increasing doses of HA-tagged MAPK6 or HA-tagged MAPK4 into the HEK293T cells. The level of the ectopically overexpressed MAPK6 was significantly lower than that of MAPK4 (both detected by anti-HA Western blots, Figure 3A), which agrees with a previous report on the rapid turnover of MAPK6 protein (20). Although expressed at lower levels, MAPK6 exhibited similar activities as MAPK4 in enhancing AKT phosphorylation at both T308 and S473 (Figure 3A). Consistently, Dox-induced MAPK6 expression significantly enhanced AKT phosphorylation and activation in PC3, DU145, OVCA433, SUM159, H1299, PNT1A, and MCF10A cells (Figure 3B). Additionally, knockdown of MAPK6 greatly inhibited AKT phosphorylation and activation in MCF7, PC3, H1299, and SUM159 cells *in vitro* and H1299 xenografts *in vivo* (Figure 3, C and D). Altogether, these data support that MAPK6 phosphorylates and activates AKT.

**Figure 3.**
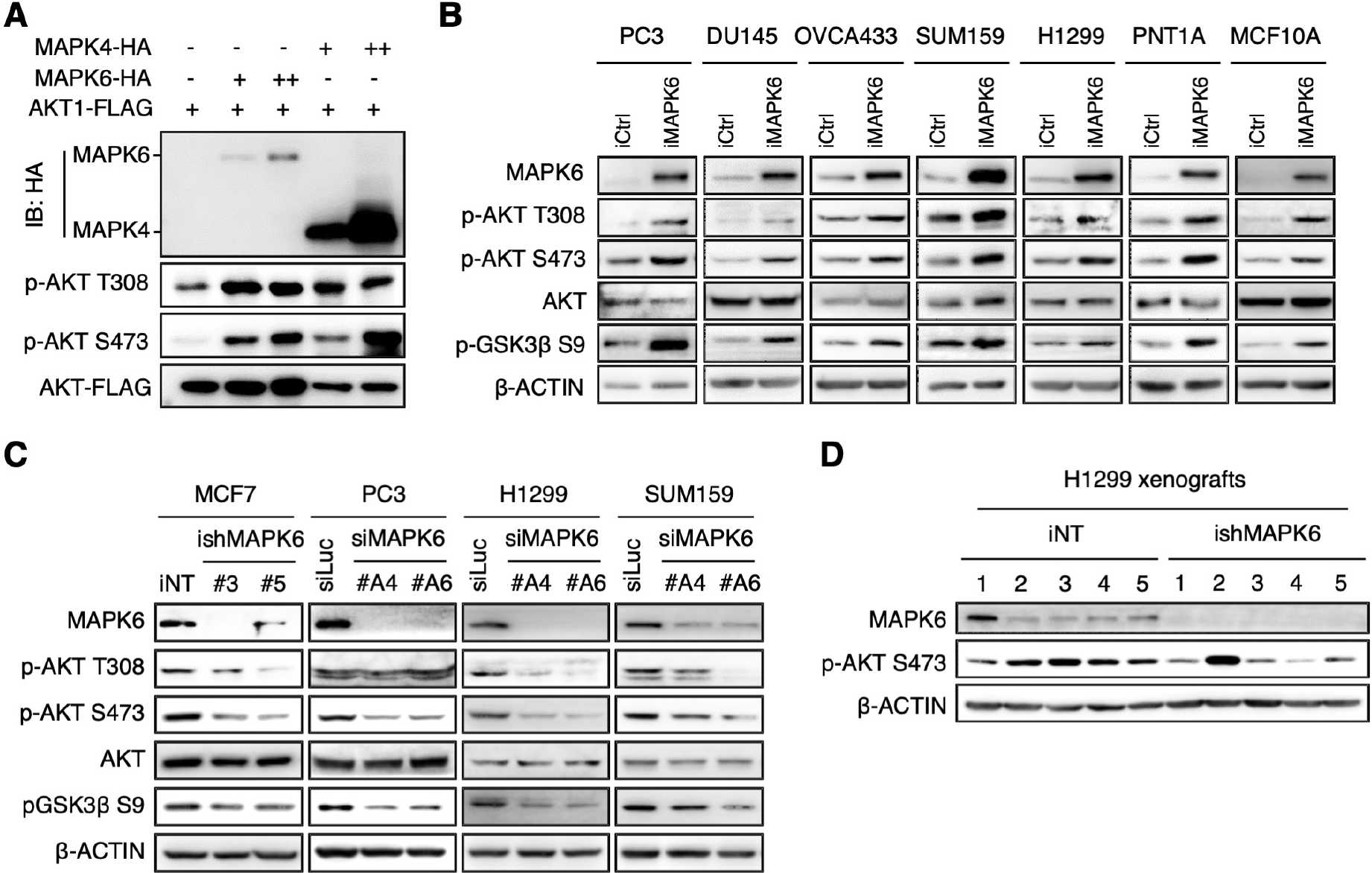
MAPK6 activates AKT. (**A**) Western blots on the HEK293T cells 48 hours after the indicated transfections. (**B**) Western blots on the Dox-induced engineered PC3, DU145, OVCA433, H1299, PNT1A, and MCF10A cells with overexpression of MAPK6 (iMAPK6) or control (iCtrl). Cells were treated with 1 μg/ml Dox for 3 to 5 days. (**C**) Western blots on MAPK6-knockdown MCF7, PC3, H1299, and SUM159 cells. The engineered MCF7 cells were treated with 4 μg/ml Dox for 5 days for knockdown of MAPK6 (ishMAPK6 #3 and #5) or control (iNT). The PC3, H1299, and SUM159 cells were transfected with siRNA against MAPK6 (siMAPK6 #A4 and #A6) or luciferase control (siLuc). 72h later, the cell lysates were prepared and applied in the Western blots. (**D**) Western blots on the H1299 xenograft tumors, as described in Figure 2E. Data are representative of at least 3 independent experiments.

### AKT activation is essential for the oncogenic and tumor-promoting activity of MAPK6

To assess the functional significance of AKT activation in mediating the oncogenic and growth-promoting activity of MAPK6, we performed soft-agar assays on Dox-induced PNT1A-iMAPK6 cells treated with AKT inhibitors MK2206, GSK2141795, or vehicle control. While MAPK6 overexpression transformed the PNT1A cells into anchorage-independent growth, MK2206 or GSK2141795 treatment largely abolished this growth (Figure 4A). Western blots confirmed that both MK2206 and GSK2141795 significantly repressed MAPK6-induced AKT activation as indicated by GSK3β phosphorylation at S9 (Figure 4B). Inhibiting AKT also blocked both basal and MAPK6-induced anchorage-independent growth of OVCA433 cells (Figure 4, C and D). Finally, overexpression of a constitutively active AKT1^T308D/S473D^ (DD) mutant rescued the anchorage-independent growth of MAPK6-knockdown H1299 cells (Figure 4, E and F). Altogether, these data strongly support an essential role for AKT activation in mediating the oncogenic and tumor-promoting activities of MAPK6.

**Figure 4.**
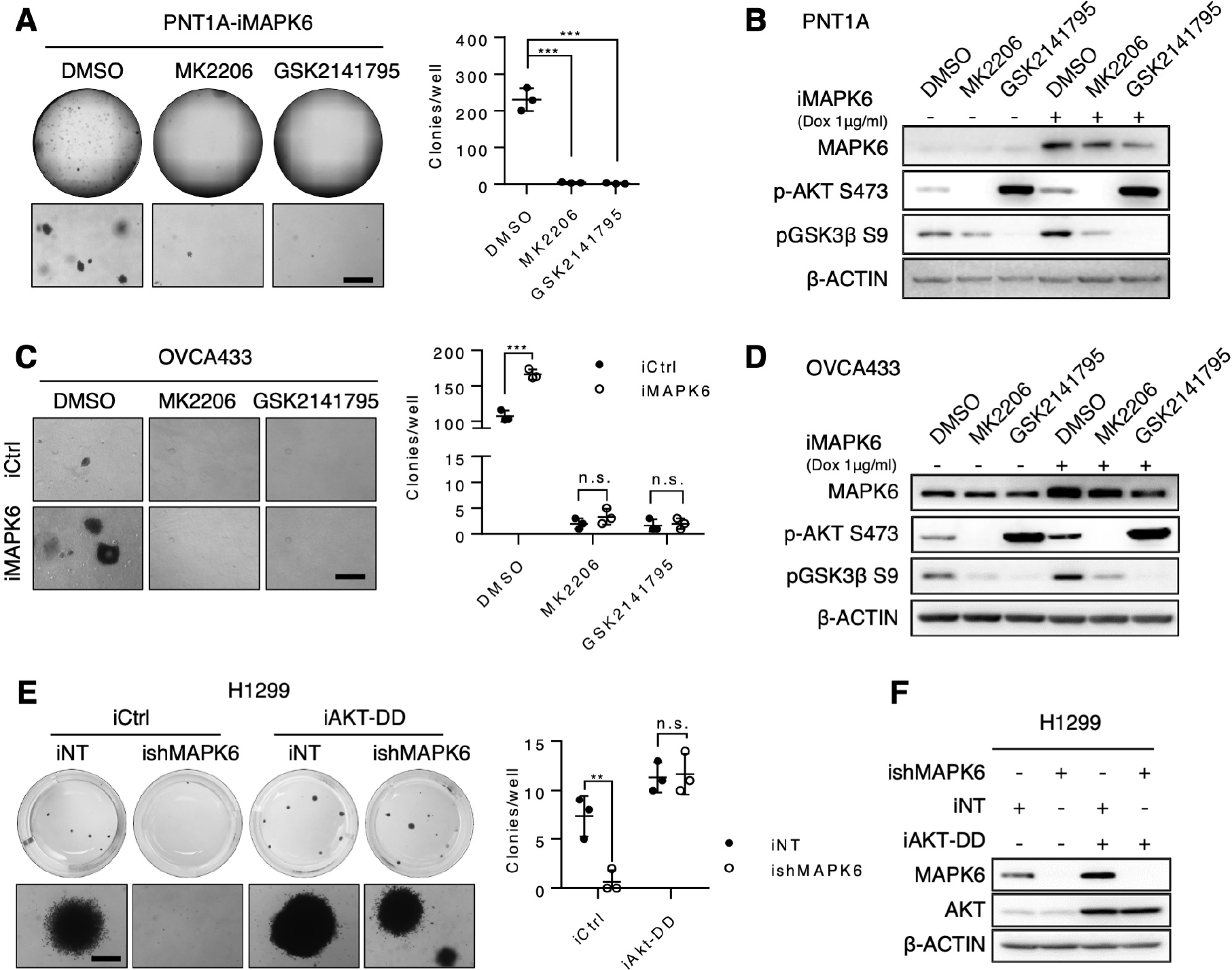
AKT activation is essential for the oncogenic and tumor-promoting activity of MAPK6. (**A**) Left panel: Representative images (whole well and enlarged field) of the soft-agar assays on 0.5 µg/ml Dox-induced PNT1A-iMAPK6 cells treated with 2 µM of MK2206, GSK2141795, or DMSO vehicle control. Right Panel: quantification of the soft-agar assay results. (**B**) Western blots on the PNT1A-iMAPK6 cells induced with 0.5 μg/ml Dox for 48h followed by treatments of 2 µM MK2206, 2 µM GSK2141795, or DMSO vehicle control for 18h. (**C** and **D**) The similar studies as described in (A) and (B), but on the engineered OVCA433 cells with 1 μg/ml Dox-induced overexpression of MAPK6 (iMAPK6) vs control (iCtrl). (**E**) Left panel: representative images (whole well and enlarged field) of the soft-agar assays on 1 µg/ml Dox-induced engineered H1299 cells with overexpression of a constitutively active AKT1^T308D/S473D^ (iAKT-DD) vs. control (iCtrl) with knockdown of MAPK6 (ishMAPK6) vs. control (iNT). Right panel: quantification of the soft-agar assay results. (**F**) Western blots on the engineered H1299 cells as described in (E). The cells were treated with 1 μg/ml Dox for 48 h and then harvested for western blots analysis. All the quantitative data were represented as mean ± SD with *P* value detected by Student’s t-test. **: *P* < 0.01. ***: *P* < 0.001. n.s.: not significant. Data are representative of at least 3 independent experiments.

### MAPK6 phosphorylates AKT at S473

The activation of AKT by MAPK6 could be either direct or indirect. To determine whether MAPK6 can directly bind and phosphorylate AKT, we affinity purified FLAG-tagged MAPK6 expressed in PC3 and HEK293T cells using anti-FLAG antibody conjugated beads. The purified MAPK6 protein contained a major band of around 110 KDa, as detected by coomassie blue staining (Figure 5A). We then performed *in vitro* kinase assays using commercially available purified AKT1 protein as substrate, as previously described (21). MAPK6 protein, purified from either the HEK293T cells or the PC3 cells, strongly phosphorylated AKT1 on S473 but not T308, identifying MAPK6 as a novel AKT S473 kinase (Figure 5B).

**Figure 5.**
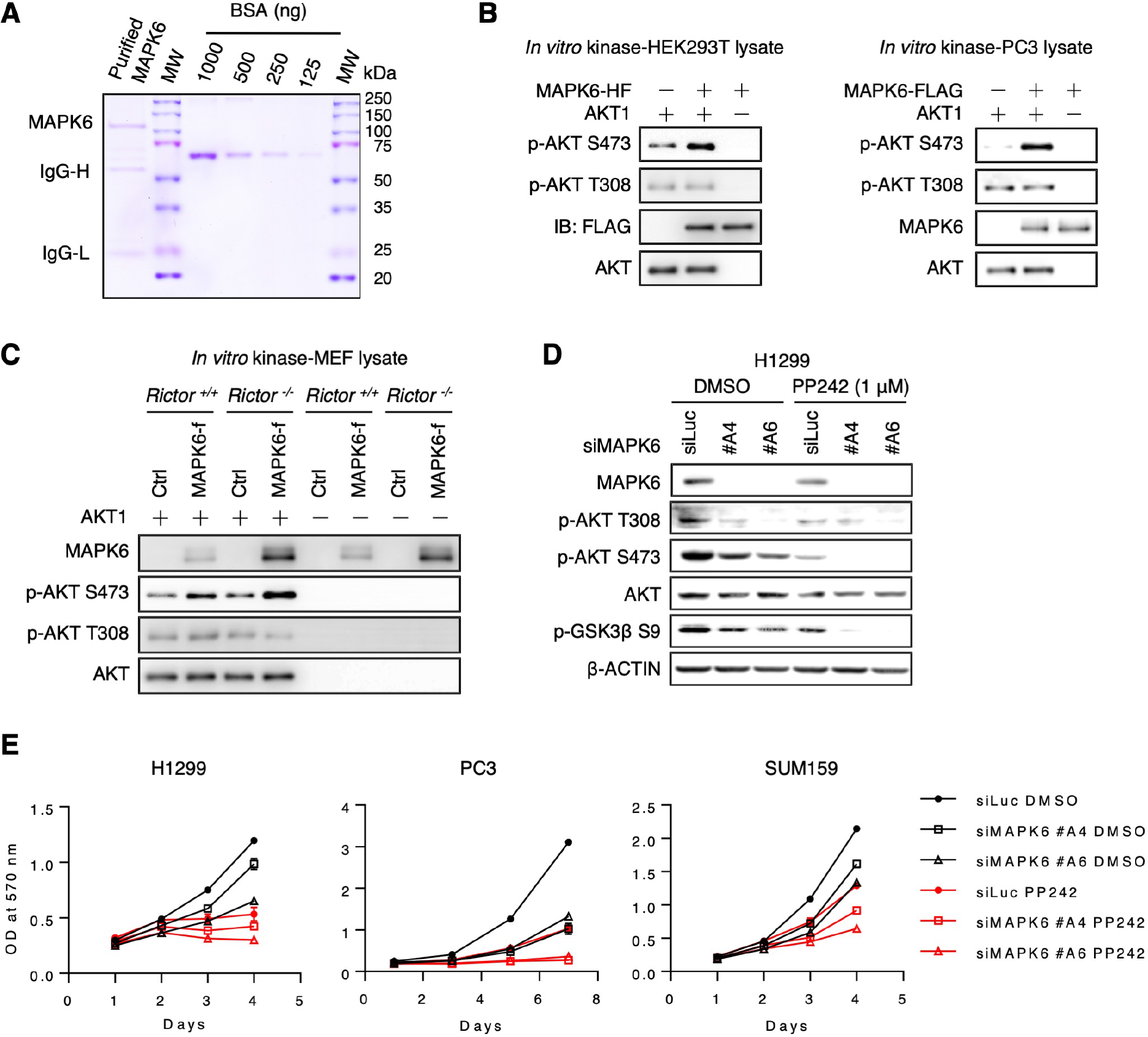
MAPK6 phosphorylates AKT at S473 independent of mTORC2. (**A**) Coomassie blue staining of the purified MAPK6 protein from HEK293T cells. MW: molecular weight. IgG-H: heavy chain of IgG. IgG-L: light chain of IgG. (**B**) Western blots on the products from the *in vitro* kinase assays assessing MAPK6 phosphorylation of AKT. MAPK6 proteins were overexpressed and purified using the EZview Red anti-FLAG M2 affinity gel from the HEK293T cells transiently transfected with the pRK5-MAPK6-HF plasmid (left panel) and the engineered PC3 cells overexpressing MAPK6-FLAG (right panel). HF: 10 x His and 2 x FLAG tag. (**C**) *In vitro* kinase assays assessing AKT phosphorylation by the MAPK6-f (MAPK6-FLAG) proteins purified from the engineered *Rictor*^*+/+*^ and *Rictor*^*−/−*^ MEF cells with 0.5 µg/ml Dox-induced expression of MAPK6-f or control (iCtrl). Shown are Western blots. (**D**) Western blots on the H1299 cells transfected with siRNA targeting MAPK6 (siMAPK6 #A4 and #A6) or control (siLuc) for 72 h followed by treatments using 1 µM PP242 or DMSO for 18h. (**E**) Proliferation assays on the H1299, PC3, and SUM159 cells with siRNA mediated knockdown of MAPK6 (siMAPK6 #A4 and #A6) or control (siLuc), treated with PP242 or DMSO. 48h after siRNA transfection, the cells were plated into 24-well plate and treated with PP242 (0.5 μM for H1299 and 1 μM for PC3 and SUM159 cells). Data are representative of at least 3 independent experiments.

mTORC2, consisting of mTOR, RICTOR, SIN1, and mLST8/GβL, is a key AKT S473 kinase (21). Our *in vitro* kinase assay data suggests mTORC2-independent S473 phosphorylation of AKT by MAPK6. To further confirm this, we expressed and affinity-purified MAPK6-FLAG proteins from *Rictor* knockout MEFs (22) and wild type MEFs using the anti-FLAG antibody conjugated beads. As expected, purified MAPK6 from *Rictor* knockout MEFs maintained its ability to phosphorylate AKT1 S473 *in vitro* (Figure 5C).

Finally, we treated MAPK6-knockdown H1299 cells and controls with PP242, an mTOR kinase-specific inhibitor (23). As expected for independent functions of the mTORC2 and MAPK6 pathways, knockdown of MAPK6 alone or PP242 treatment alone significantly reduced AKT S473 phosphorylation, but co-treatment largely abolished it (Figure 5D). In accord with this, the combination of MAPK6 knockdown and PP242 treatment much more strongly suppressed H1299, PC3, and SUM159 cell proliferation than either treatment alone (Figure 5E). Collectively, these results support MAPK6 phosphorylation of S473 independent of mTORC2.

### MAPK6 directly binds AKT

To further define the MAPK6-AKT pathway, we investigated MAPK6-AKT interaction. We expressed FLAG-tagged AKT1 in HEK293T cells overexpressing HA-tagged MAPK6 and performed immunoprecipitation using anti-FLAG antibody conjugated beads followed by Western blot for HA. We also expressed FLAG-tagged MAPK6 in HEK293T cells overexpressing a His-tagged AKT1 and performed immunoprecipitation using anti-FLAG antibody conjugated beads followed by Western blot for AKT. In both cases, we readily detected MAPK6 and AKT1 interaction (Figure 6A).

**Figure 6.**
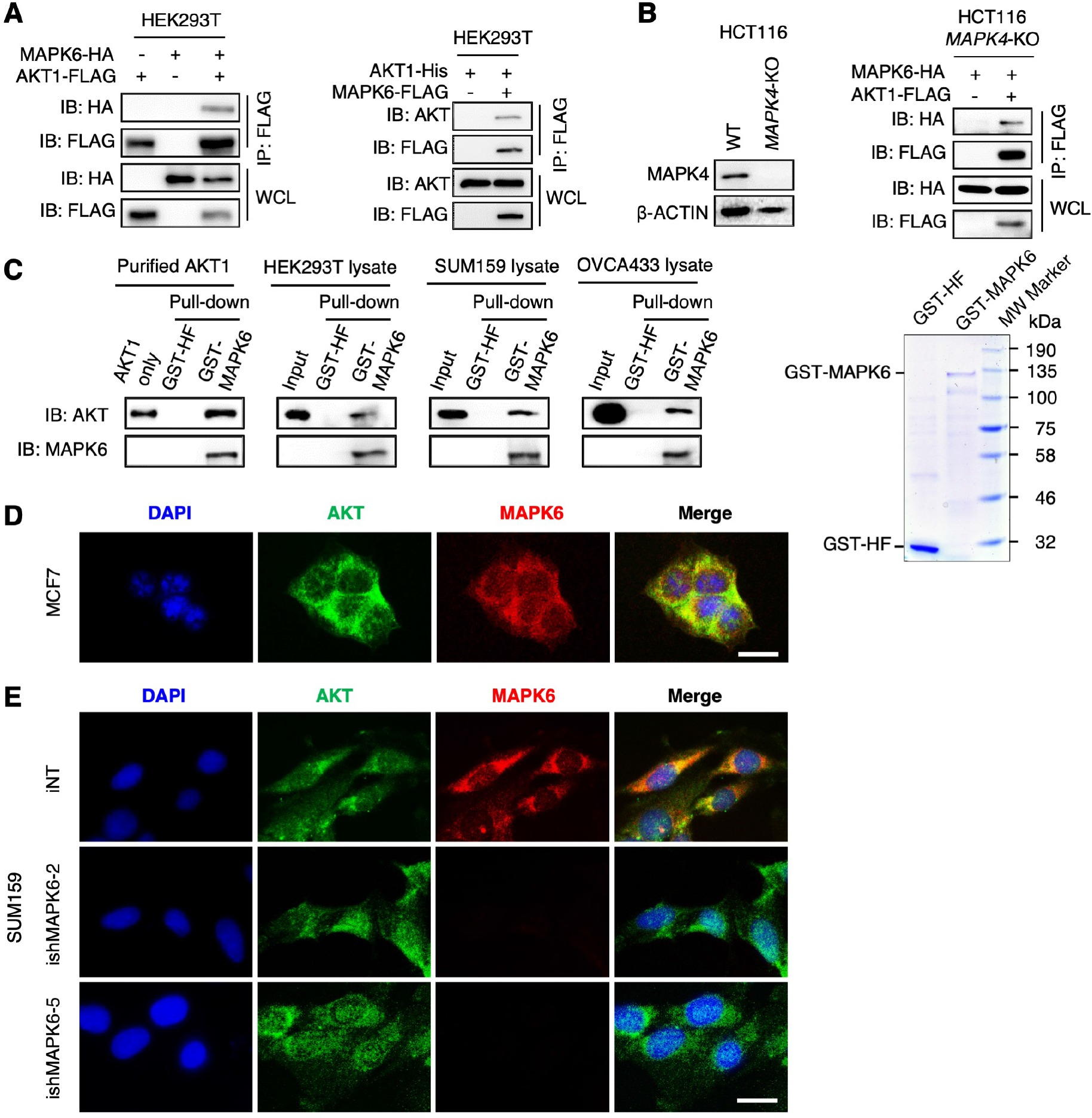
MAPK6 binds to AKT. (**A**) Western blots on the immunoprecipitation products using anti-FLAG M2 affinity gel. HEK293T cells were transfected with MAPK6-HA and AKT1-FLAG (left Panel) or AKT1-His and MAPK6-FLAG (Right Panel). 48h later, the cell lysates were prepared for the immunoprecipitation using anti-FLAG M2 affinity gel and Western blots. (**B**) Left Panel: Western blots on MAPK4 expression in the wild type and *MAPK4*-KO HCT116 cells. Right Panel: Western blots showing the co-IP of MAPK6-HA and AKT1-FLAG in the absence of MAPK4. The *MAPK4*-KO HCT116 cells were transfected with MAPK6-HA and AKT1-FLAG. 48h later, the cell lysates were prepared for immunoprecipitation using anti-FLAG M2 affinity gel and Western blots. (**C**) GST pull-down assay showing binding between purified GST-MAPK6 and purified AKT1 as well as endogenous AKT in various cell lysates. Coomassie blue staining (right panel) revealed a major band of around 130 kDa and 30 kDa in the purified GST-MAPK6 and GST-HF, respectively. MW: molecular weight. HF: 10x His and 2x FLAG tag. (**D**) Immunofluorescence staining showing the co-localization of endogenous MAPK6 and AKT in the cytoplasm of MCF7 cells. Label: 20 μm. (**E**) Immunofluorescence staining showing the co-localization of endogenous MAPK6 and AKT in the cytoplasm of SUM159 cells. The specificity of MAPK6 antibody was confirmed by a lack of immunofluorescence signal in the MAPK6-knockdown (ishMAPK6-2 and ishMAPK6-5) cells. Label: 20 μm. Data are representative of at least 3 independent experiments.

MAPK6 and MAPK4 can form a heterodimer (24), and we reported that MAPK4 could bind AKT (19), raising the prospect that MAPK4 could mediate apparent MAPK6-AKT interaction. To critically test this, we expressed FLAG-tagged AKT1 in *MAPK4*-KO HCT116 cells (19) overexpressing HA-tagged MAPK6 and performed immunoprecipitation using anti-FLAG antibody conjugated beads followed by Western blot for HA. Again, we confirmed MAPK6-AKT interaction in the absence of MAPK4 (Figure 6B). The direct interaction of MAPK6 and AKT was strongly confirmed by the demonstration that overexpressed and purified GST-MAPK6 fusion protein, but not GST alone, can pull down commercially available purified AKT1 protein (Figure 6C). Pull-down assays using HEK293T, SUM159, and OVCA433 cell lysates confirmed that GST-MAPK6 protein, but not GST, can pull down the endogenous AKT in these cell lysates (Figure 6C).

MAPK6 protein locates in both cytoplasm and nucleus (16). Since AKT primarily acts in the cytoplasm, we expected colocalization of cytoplasmic MAPK6 and AKT. Immunofluorescence using anti-MAPK6 and anti-AKT antibodies confirmed such cytoplasmic colocalization in MCF7 (Figure 6D) and SUM159 cells (Figure 6E). MAPK6 knockdown abolished this co-immunofluorescence (Figure 6E). These data support the notion that MAPK6 binds to AKT in the cytoplasm for MAPK6-induced AKT activation.

### MAPK6 and MAPK4 bind AKT by different mechanisms

MAPK6 contains an N-terminal kinase domain, a C34 conserved region shared between MAPK6 and MAPK4, and a unique long C-terminal tail (Figure 7A). Since we previously identified a new D/E-enriched common docking (CD) motif within MAPK4 (EEDKDE at positions 250-255) providing the critical docking site for AKT binding, we first investigated whether AKT binds to MAPK6 via the similar D/E-enriched motif (EEDRQE) at positions 253-258 of MAPK6 (Figure 7B, upper panel). For this purpose, we expressed a FLAG-tagged MAPK6 mutant with EEDRQE mutated to AAARAA (D/E/Q-5A) and His-tagged AKT1 in HEK293T cells and performed immunoprecipitation using anti-FLAG antibody conjugated beads followed by Western blot for AKT. Mutation of these D/E/Q residues had minimal effect on MAPK6-AKT interaction (Figure 7B), indicating that this putative CD motif is not required for MAPK6 binding to AKT.

**Figure 7.**
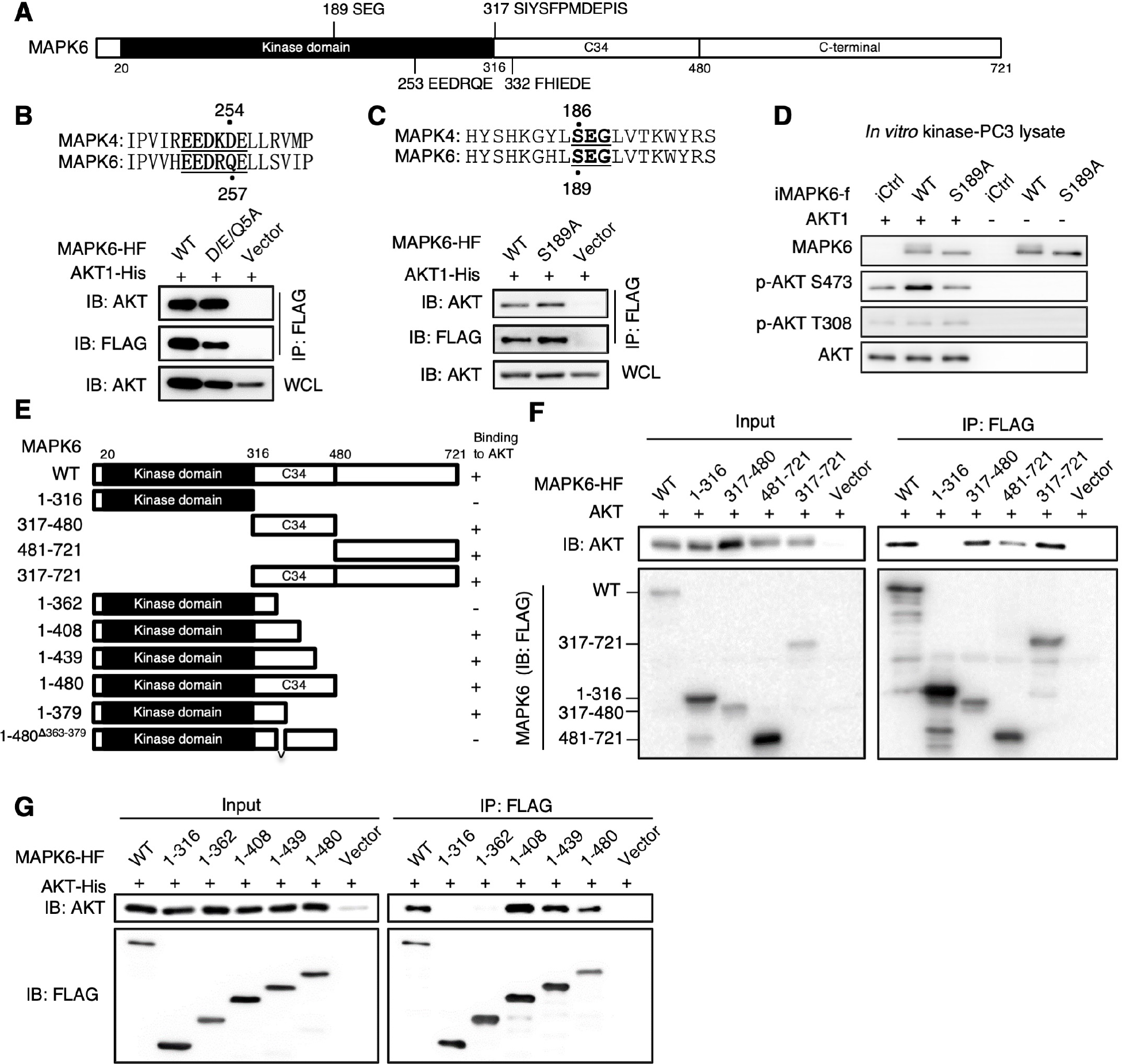
The molecular basis for MAPK6 binding to AKT, part 1. (**A**) Diagram showing the domain architecture of human MAPK6. MAPK6 contains an N-terminal domain (20 aa to 316 aa), a conserved C34 (317 aa to 480 aa) shared between MAPK6 and MAPK4, and a C-terminal tail (481 aa to 721 aa). (**B**) Western blots on the IP products for interaction between AKT1-his and wild type or a mutant (D/E/Q5A: EEDRQE to AAARAA) MAPK6. HEK293T cells were transfected with AKT1-His along with wild type (WT) or a MAPK6 mutant (D/E/Q5A). 48h later, the cell lysates were prepared for immunoprecipitation using anti-FLAG M2 affinity gel, followed by Western blots. WCL: whole cell lysate. (**C**) Western blots on the IP products for interaction between AKT1-His and wild type (WT) or a mutant (S189A) MAPK6. The HEK293T cells were similarly transfected and processed as described in (B). (**D**) *In vitro* kinase assays comparing wild type (WT) and S189A mutant MAPK6 in phosphorylating AKT1. *In vitro* kinase assays were carried out using WT and S189A mutant MAPK6 proteins expressed and purified from the engineered PC3 cells as similarly described in Figure 5A. (**E**) Diagram showing the fragments and mutants of MAPK6 used in (F), (G), (H), and Figure 8 C and D. (**F-G**) Western blots on the input and IP products for interaction between AKT1 and truncated MAPK6. The HEK293T cells were similarly transfected and processed, as described in (B). Wild Type (WT) MAPK6 was used as positive control. (F) AKT was bound to both C34 and C-terminal tail of MAPK6, but not the kinase domain. (G) A fragment between 362 aa to 408 aa was essential for MAPK6 to bind to AKT. aa: amino acid. Data are representative of at least 3 independent experiments.

Conventional MAP kinases share a conserved TXY motif in the activation loop that is the target for MAPK kinase (MAPKK) induced serine/threonine and tyrosine dual phosphorylation. In MAPK4 and MAPK6, this is replaced by SEG (S186 for MAPK4 and S189 for MAPK6, Figure 7C, upper panel). We have shown that S186A mutation inhibited MAPK4 binding and phosphorylation of AKT (19). To assess whether S189 also plays a similar role in MAPK6, we introduced His-tagged AKT1 and FLAG-tagged wild type MAPK6 or the S189A mutant MAPK6 (MAPK6^S189A^) into HEK293T cells. We found that unlike the MAPK4^S186A^ mutant, the MAPK6^S189A^ mutant maintained AKT1 interaction (Figure 7C). However, the MAPK6^S189A^ mutant exhibited little activity in phosphorylating AKT1 S473 in the *in vitro* kinase assay (Figure 7D).

Altogether, our data strongly suggest that MAPK6 uses a different mechanism to bind to AKT. This MAPK6-AKT binding is neither dependent on the previously identified D/E-enriched CD motif nor dependent on the SEG motif.

To determine the critical region of MAPK6 required for AKT interaction, we performed co-immunoprecipitation of His-tagged AKT1 with FLAG/His-tagged fragments of MAPK6 overexpressed in HEK293T cells. In contrast to the MAPK4 kinase domain, which can efficiently bind AKT (19), the kinase domain of MAPK6 (the 1-316 aa fragment of MAPK6, MAPK6^1-316^) failed to interact with AKT (Figure 7 E and F). Instead, the C34 region (the 317-480 aa fragment of MAPK6, MAPK6^317-480^), the C-terminal tail (MAPK6^481-721^), and the fragment containing both C34 and C-terminal tail (MAPK6^317-721^) were sufficient to bind AKT (Figure 7, E and F). The C34 region appeared to exhibit a higher AKT binding affinity than that of the C-terminal tail (Figure 7, E and F). These data suggest that MAPK6 binds to AKT through both the higher affinity C34 region and the lower affinity C-terminal tail.

To map the high affinity site, we further created a series of C-terminally truncated MAPK6 containing either the intact or part of the C34 region (Figure 7E). MAPK6 fragments MAPK6^1-480^, MAPK6^1-439^, and MAPK6^1-408^, but not MAPK6^1-362^ or MAPK6^1-316^, maintained high affinity to AKT (Figure 7G). This suggests that the 362-408 region of MAPK6 contains a motif critically required for MAPK6-AKT interaction. To further map and define the critical region for MAPK6-AKT binding, we also generated additional C-terminally truncated fragments. MAPK6^1-379^, MAPK6^1-384^, and MAPK6^1-389^, but not MAPK6^1-373^ or MAPK6^1-362^, maintained their affinity to AKT (Figure 8A), identifying the 374-379 aa region of MAPK6 containing at least part of the sequence critical for AKT binding. In accord with their distinct AKT binding properties, MAPK6 and MAPK4 share little homology within this 374-379 or the broader 363-379 region (Figure 8B and Supplementary Figure S2, A and B).

**Figure 8.**
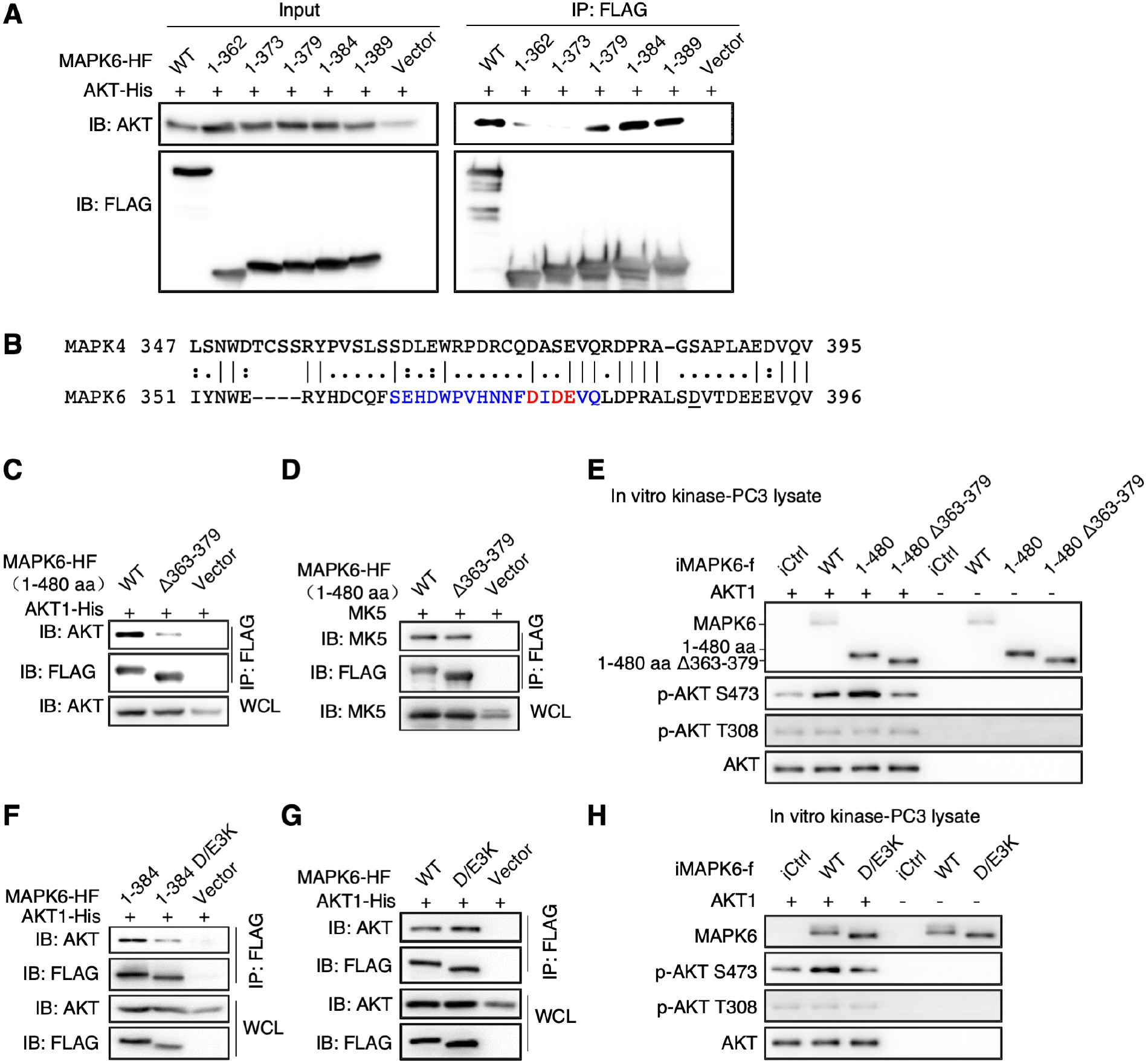
The molecular basis for MAPK6 binding to AKT, part 2. (**A**) Western blots on the input and IP products for interaction between AKT1 and truncated MAPK6. The HEK293T cells were similarly transfected and processed, as described in Figure 7B. MAPK6^1-379^ but not MAPK6^1-373^ was efficient to bind to AKT. (**B**) Sequence alignment between MAPK6 and MAPK4 around the potential AKT binding motif of MAPK6. (**C** and **D**) Western blots on the IP products for interaction between AKT1 and truncated MAPK6. Deletion of 363-379 aa of MAPK6 (Δ363-379, SEHDWPVHNNFDIDEVQ labeled in blue/Red in (B)) suppressed MAPK6^1-480^ interaction with AKT1 (C) but not MK5 (D). The HEK293T cells were similarly transfected and processed, as described in Figure 7B. Wild Type (WT) MAPK6 was used as positive control. (**E**) *In vitro* kinase assays comparing wild type (WT) MAPK6, MAPK6^1-480^ and MAPK6^1-480 Δ363-379^ ability in phosphorylating AKT1. Assays were carried out using the above MAPK6 proteins expressed and purified from the engineered PC3 cells, as similarly described in Figure 5A. (**F** and **G**) Western blots on the IP products for interaction between AKT1 and the MAPK6 mutants. The mutation of DIDE (374-377) highlighted in red in (B) into KIKK (D/E3K) blocked MAPK6^1-384^ (F) but not full-length MAPK6 (G) binding to AKT1. (**H**) *In vitro* kinase assays comparing wild type (WT) and the D/E3K mutant MAPK6 ability in phosphorylating AKT1. Assays were carried out using the above MAPK6 proteins expressed and purified from the engineered PC3 cells, as similarly described in Figure 5A. WCL: whole cell lysate. HF: 10 x His and 2 x FLAG tags. See also Figure S2. Data are representative of at least 3 independent experiments.

Accordingly, the MAPK6^1-480^ mutant with an internal 363-379 deletion (MAPK6^1-480 **Δ**363-379^) mostly lost AKT binding (Figure 8C) despite maintaining MK5 binding through the FHIEDE motif (Figure 8D and Figure 7A). Accordingly, MAPK6^1-480^ exhibited kinase activity in phosphorylating AKT at S473 in the *in vitro* kinase assay, while MAPK6^1-480 Δ363-379^ largely lost this activity (Figure 8E). Altogether, these data suggest that the 363-379 region of MAPK6 is critical for MAPK6-AKT interaction.

We hypothesized that a candidate D/E-enriched motif within the 363-379 aa region of MAPK6 (DIDE, 374-377 aa) might be a CD motif for MAPK6 and AKT interaction. In accord with this, mutation of DIDE to KIKK (D/E3K) abolished binding between MAPK6^1-384^ and AKT (Figure 8F). However, this D/E3K mutation did not affect binding between full-length MAPK6 and AKT (Figure 8G), potentially due to the interaction between the MAPK6 C-terminal tail and AKT (Figure 7F). Despite the lack of impact of the D/E3K mutation on binding between full-length MAPK6 and AKT, the MAPK6^D/E3K^ mutant largely lost its ability to phosphorylate AKT at S473 in the *in vitro* kinase assay (Figure 8H). These data suggest that the binding site within the C34 region, along with the second site within the C-terminal tail, is essential for functional docking of AKT onto MAPK6 for phosphorylation/activation.

### MAPK6 overexpression correlates with poor survival in human cancers

Our data support an oncogenic and tumor-promoting activity of MAPK6 that predicts a negative correlation between *MAPK6* mRNA expression and cancer patient survival. A comprehensive analysis of gene expression profiles of 10,152 patients in The Cancer Genome Atlas (TCGA) (25) revealed that *MAPK6* is expressed at varying levels in all cancer types (Figure 9A). In this large pan-cancer panel, survival was significantly decreased in the subset of patients above the 80^th^ percentile of *MAPK6* expression (Figure 9B). Although shorter follow-up data and small numbers of patients limited analysis in some cases, survival was also markedly decreased in patients with *MAPK6* overexpression above the 75^th^ percentile in several specific cancer types in the TCGA dataset, including lung adenocarcinoma (LUAD), mesothelioma (MESO), and uveal melanoma (UVM). Analysis of the cancer compendium also showed decreased survival in lung adenocarcinoma (26) and breast cancer patients (27) with *MAPK6* overexpression (Figure 9C).

**Figure 9.**
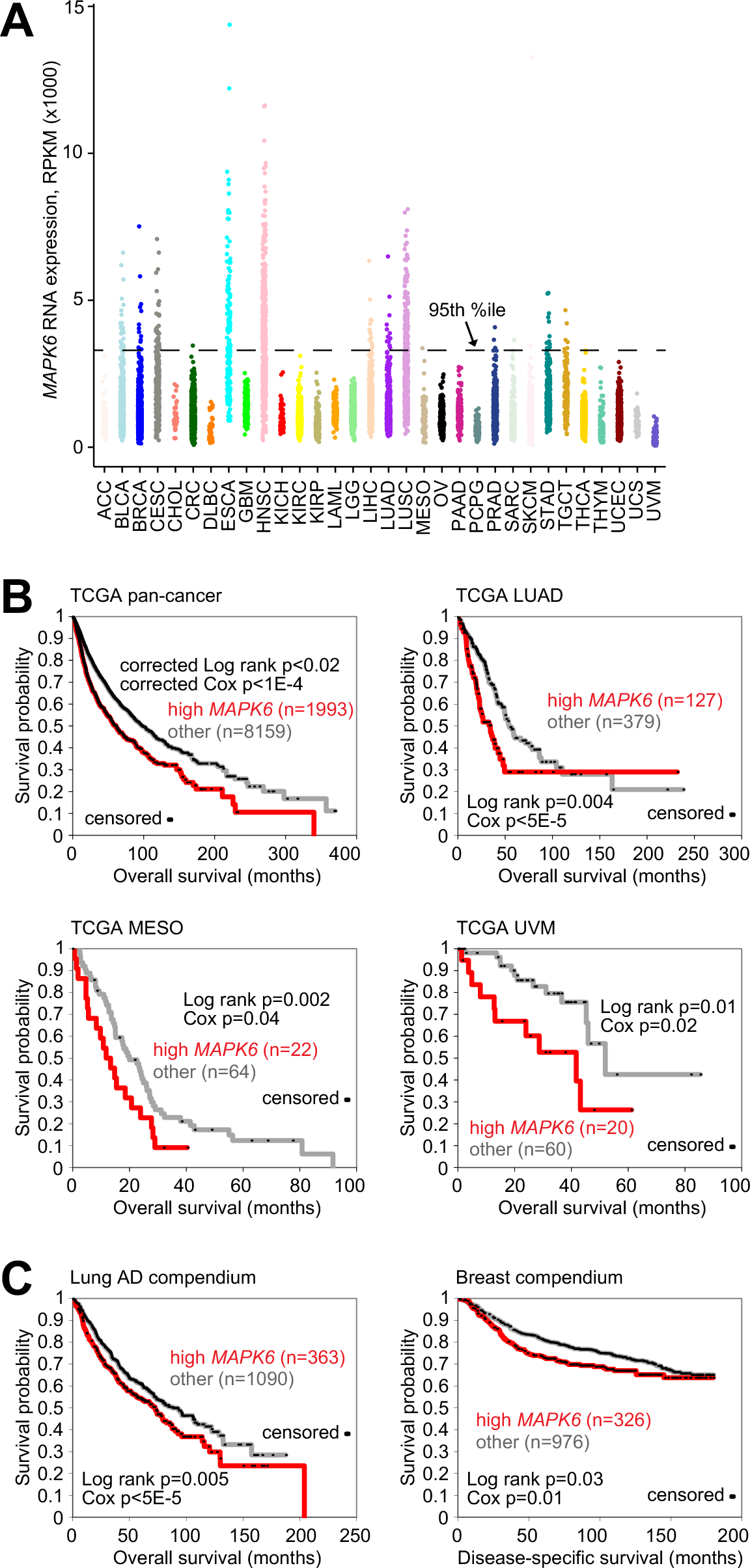
Overexpression of MAPK6 in a subset of human cancers is associated with decreased overall survival. (**A**) MAPK6 mRNA expression across 10,152 tumors of various histological subtypes from The Cancer Genome Atlas (TCGA). (**B**) Using TCGA data, Kaplan-Meier plots of overall survival in patients, both across all cancer types in TCGA cohort (“pan-cancer” analysis) or within specific cancer types as indicated, as stratified by high MAPK6 expression. *P*-values by univariate Cox or Log-rank test, as indicated. For pan-cancer plot, *P*-values by stratified log-rank test and stratified univariate Cox, each test correcting for overall differences in survival according to tumor type. (**C**) In both a compendium of lung adenocarcinoma expression profiles (left) and a compendium of breast cancer expression profiles (right), Kaplan-Meier plots of overall survival in patients, as stratified by high MAPK6 expression. *P* values by univariate Cox or Log-rank test, as indicated.

## Discussion

The limited numbers of previous studies on MAPK6 biology in human cancers have focused on regulation of the cancer cell migration/invasion/metastasis (8, 10-15). MAPK6 also promoted cervical cancer cell growth, but inhibited melanoma and intrahepatic cholangiocarcinoma cell proliferation (10, 13, 14, 17, 18). To date, it appears that whether MAPK6 executes tumor promoting or tumor suppressing function is cancer type or cancer cell context dependent. To critically examine how MAPK6 regulates human cancers, we used six common human cancer cell lines representing four different cancer types, including prostate cancer (PC3 and DU145), breast cancer (MCF7 and SUM159), ovarian cancer (OVCA433), and non-small cell lung cancer (H1299). We also assessed the oncogenic activities of MAPK6 using two non-transformed “normal” human epithelial cell lines, the prostate epithelial cell line PNT1A and the breast epithelial cell line MCF10A. We demonstrated that MAPK6 overexpression transforms both human epithelial cell lines into anchorage-independent growth and also promotes growth, including anchorage-independent growth of all five cancer cell lines tested (PC3, DU145, SUM159, H1299, and OVCA433). These data clearly define the potent oncogenic and tumor promoting activities of ectopically overexpressed MAPK6 in all of the seven cell lines examined. Accordingly, knockdown of MAPK6 in four cancer cell lines with medium to high expression of endogenous MAPK6 (MCF7 and SUM159 for breast cancer, PC3 for prostate cancer, and H1299 for non-small cell lung cancer) inhibited cell growth *in vitro* and/or xenograft growth *in vivo*.

We recently reported that MAPK4, another atypical MAPK most closely related to MAPK6, promotes cancer by non-canonically activating the key oncogenic kinase AKT independent of PI3K/PDK1 (19). MAPK4 acts as a T308 kinase and also induces mTORC2-dependent S473 phosphorylation. In contrast, MAPK6 mainly functions as an S473 kinase in the *in vitro* kinase assay, independent of mTORC2 (Figure 5, B and C). MAPK6 also induces AKT phosphorylation at T308 in cells, indicating the potential involvement of PI3K/PDK1 and/or MAPK4 for this phosphorylation. When ectopically overexpressed in HEK293T cells, MAPK6 appeared to have more robust activities than MAPK4 in phosphorylating AKT -- the HA-tagged MAPK6 exhibited similar activities in phosphorylating AKT when expressed at a significantly lower level than that of the HA-tagged MAPK4. Ectopic overexpression of MAPK6 in eight tested cell lines, including PC3, DU145, SUM159, OVCA433, H1299, PNT1A, MCF10A, and HEK293T cells, induced AKT phosphorylation/activation. In accord with this, knockdown of endogenous MAPK6 repressed AKT phosphorylation and activation in the four tested cell lines (MCF7, PC3, SUM159, and H1299) with high to median endogenous MAPK6 expression, further confirming the MAPK6-AKT pathway in the cells. Finally, knockdown of MAPK6 greatly inhibited AKT phosphorylation/activation in the H1299 xenograft tumors and repressed their growth, demonstrating the tumor-promoting MAPK6-AKT pathway *in vivo*.

Human MAPK4 and MAPK6 share a similar domain structure (1). We have previously reported that a novel CD motif within the kinase domain of MAPK4 mediates MAPK4-AKT binding, and the D254 within the CD motif is essential for this interaction. Besides, phosphorylation of S186 of the SEG motif also appears to play a critical role in regulating MAPK4-AKT interaction. Interestingly, D254 of MAPK4 is replaced by Q257 in the corresponding potential CD motif of MAPK6, suggesting that MAPK6 may use a different mechanism for AKT binding. Indeed, neither disruption of the potential CD motif around Q257 nor mutation of S189 (corresponding to S186 in MAPK4) affected MAPK6-AKT interaction. In accord with this, the kinase domain of MAPK6, unlike that of MAPK4, lacks binding affinity to AKT. Other than the D254/Q257 variation, the CD motifs of MAPK4 and MAPK6 within the kinase domain are highly conserved. Interestingly, a Q257D mutation largely restored MAPK6 kinase domain binding with AKT, confirming the potential functionality of this largely conserved CD motif in MAPK6 (data not shown). This further confirms, in a different context, the essential role of D254 in MAPK4-AKT binding.

MAPK4 and MAPK6 share relatively high homology within the C34 region. We identified that the SEHDWPVHNNFDIDEVQ fragment at 363 to 379 aa of the MAPK6 C34 region is critical for MAPK6^1-480^ binding to AKT. Interestingly, this sequence is not conserved in MAPK4, which may underlie its unique ability to bind AKT. The unique long C-terminal tail of MAPK6 also contains an AKT binding activity with somewhat lower apparent affinity. All in all, in contrast to the MAPK4 kinase domain mediated binding of AKT, MAPK6 uses a distinct mechanism based on unique sequences within C34 and the C-terminal tail.

AKT activity was required for MAPK6 oncogenic and tumor-promoting activities under the conditions that we examined. Since both the canonical PI3K/PDK1 pathway and our recently identified MAPK4 pathway can independently regulate AKT activation, however, the tumor-promoting activities of endogenous MAPK6 are likely to be context-dependent. Since MAPK6 has been reported in both the cytoplasm and the nucleus, it is expected that activation of cytoplasmic AKT will not be significant in cancer cells where MAPK6 mainly locates in the nucleus. Our results highlight the importance of understanding the molecular mechanism underlying MAPK6 cytoplasm-nucleus translocation and its functional significance.

The pro-oncogenic role of MAPK6 described here is strongly supported by the observation that elevated *MAPK6* mRNA expression negatively correlates with pan-cancer patient survival and survival in specific cancer types, including lung adenocarcinoma, mesothelioma, uveal melanoma, and breast cancer. This raises the prospect that blocking MAPK6, either alone or in combination with mTOR inhibitors (Figure 5, D and E), will be an effective therapeutic for MAPK6-high cancers.

## Methods

### Plasmids

The pInducer20-YF vector was generated as previously described [2]. pRK5 vector was provided by Dr. Xin-Hua Feng at Baylor College of Medicine, Houston, Texas. The CDS of *MAPK6* was PCR amplified from the LNCaP cell cDNA and sequencing verified to contain the same sequence as CDS in NM_002748 in Gene Bank. The CDS of *MAPK6* was cloned into the pInducer20-YF vector between the MluI and SalI sites. The full-length, truncated, and mutated CDS of *MAPK6* were also cloned into the pRK5 vector between the MIuI and PacI sites. CDS of 1-480 aa of MAPK6 was cloned into pInducer20-YF between the MluI and XhoI sites from the pRK5-MAPK6^1-480^-HF. The same strategy was used for generating pInducer-MAPK6^1-480 Δ363-379^-HF. AKT1^K385A^, AKT1^K386A^, and AKT1^K389A^ were reported previously (19). Primers used for generating AKT1^R391A^ are as follows: R: CAAGAAGGACCCCAAGCAGGCGCTTGG and F: CCAAGCGCCTGCTTGGGGTCCTTCTTG. Primers used for cloning of wild type, mutant, and truncated MAPK6 are as described in Table I.

**Table I.**
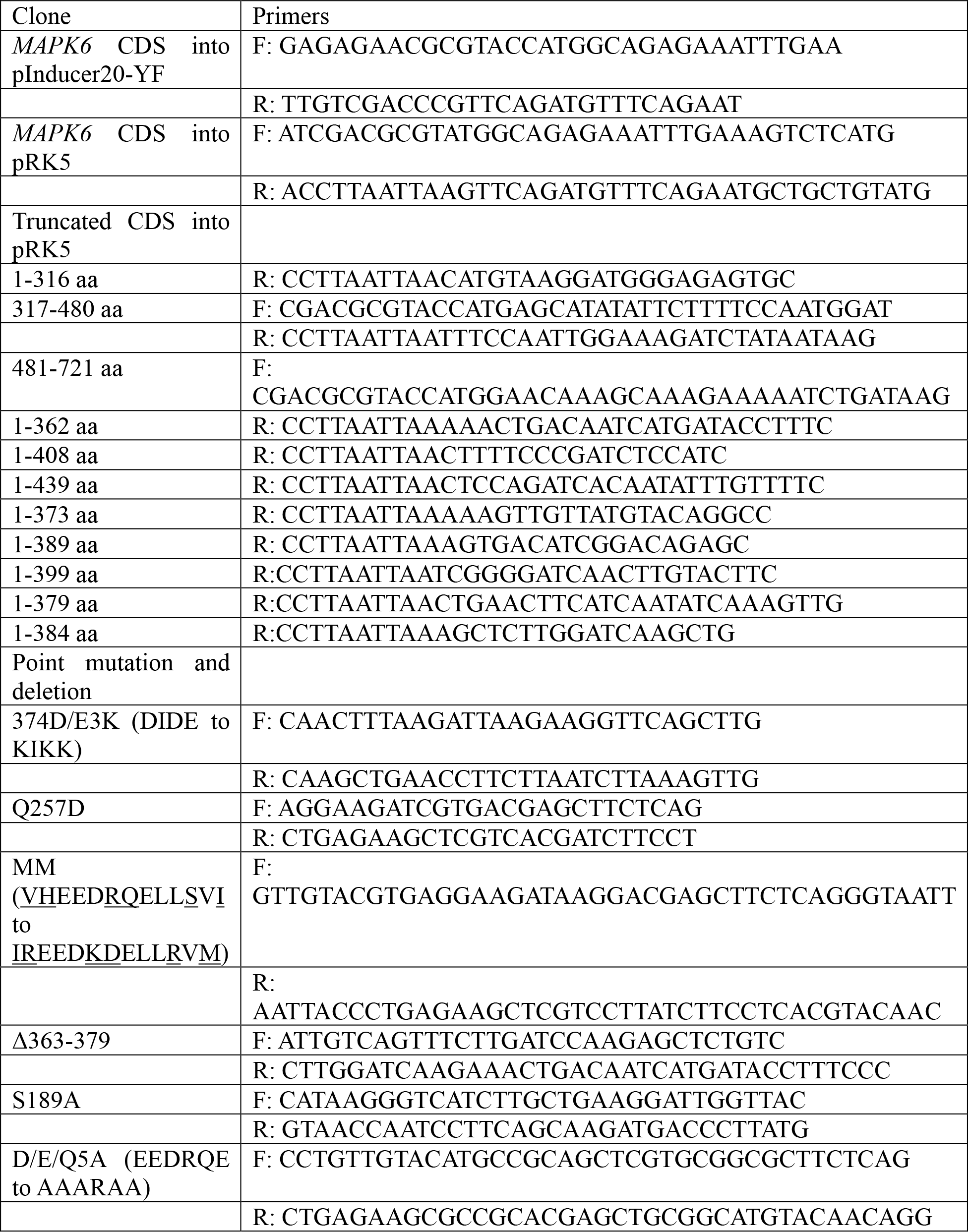
Primers used in this study.

### Cell culture, transient transfection, and lentivirus packaging

MEF *Rictor*^*+/+*^ and MEF *Rictor*^*–/–*^ cells were obtained from Mark Magnuson at Vanderbilt University, Nashville, Tennessee (22). HCT116 *MAPK4*^−/−^ cells were generated in our lab as previously described (19). PNT1A cells were acquired from the European Collection of Authenticated Cell Cultures (ECACC). Other cell lines were obtained from the American Type Culture Collection (ATCC). MCF10A cells were cultured in DMEM medium supplemented with 5% horse serum (Gibico #16050122), 0.5 μg/ml hydrocortisone (Sigma #H-0888), 20 ng/ml hEGF (Sigma #E9644), 10 μg/ml insulin (Santa Cruz #CAS11061-68-0), 100 ng/ml cholera toxin (Sigma C-8052), 100 units/ml penicillin, and 100 μg/ml streptomycin (GenDEPOT #CA005). MCF7 cells were cultured in DMEM medium with 10 μg/ml insulin, HCT116, PNT1A and PC3 cells in RPMI 1640, and other cell lines in DMEM, all supplemented with 10% fetal bovine serum, 100 units/ml penicillin, and 100 μg/ml streptomycin. Transient transfection and lentivirus packaging were done as we previously reported (19).

### shRNAs and siRNAs for knockdown

Lentiviral shRNA constructs were obtained from Open Biosystem (Thermo Fisher Scientific). Targeting sequences were cut from the pGIPZ constructs using MIuI and XhoI sites and cloned into pInducer10 using the same sites. The targeting sequences are shown as follows: pGIPZ-shMAPK6-#2: AGAGTGTCAAACATGCTCT, pGIPZ-shMAPK6-#3: GTCATCATCATATTTAGAT, and pGIPZ-shMAPK6-#5: TGGGTCACCACTTAAGTCA. siRNAs were obtained from Invitrogen as following, siMAPK6-#A4: GAGCUCUGUCCGAUGUCACUGAUGA and siMAPK6-#A6: GCCUUGUUGGCAAUACUCAGAUCAU.

### Proliferation assay

Cells were seeded at 5,000-20,000 cells per well (cell line dependent) in 500 μl media in a 24-well plate. Cell culture was stopped at the predetermined time points and then fixed with 10% (w/v) formaldehyde for 15 min and stained with 0.05% (w/v) crystal violet supplemented with 10% (v/v) ethanol and 10% (v/v) methanol for 20 min at room temperature. After washing and drying, 300 μl 10% acetic acid was added to each well and absorbance was read at 570 nm. Doxycycline was used for induced gene overexpression (1 μg/ml) or knockdown (4 μg/ml). For proliferation assays using cells with siRNA-mediated knockdown of MAPK6, the cells were first transfected with 10 nM siRNA for 48 h and then plated in 24-well plate for proliferation assays. When applicable, mTOR kinase inhibitor PP242 (Millipore Sigma P0037) was added to cell culture media the day after cell seeding.

### Plate colony formation assay and soft agar assay

For plate colony assay, 500-1000 cells/well (cell line dependent) were seeded in 6-well plates in 2 ml media and replaced with new media every week. After 2-3 weeks, cells were fixed with 10% (w/v) formaldehyde for 15min and stained with 0.05% (w/v) crystal violet supplemented with 10% ethanol and 10% methanol for 20 min at room temperature. The staining solution was then removed and wells washed with distilled water. The plates were then air-dried and scanned using a Canon scanner, and colony numbers were calculated. Soft agar assays were performed as previously described (19). AKT inhibitors MK2206 (Selleck #S1078) and GSK1411795 (Selleck #S7492) were added during setup and replaced with fresh media every week.

### Western blotting

Protein samples were prepared in RIPA buffer (100 mM NaCl, 0.5% sodium deoxycholate, 0.1% SDS, 50 mM Tris-HCl, pH 8.0, 1% Triton X-100 with 100× protease inhibitors cocktail solution from GenDEPOT [P3100-005] and the phosphatase inhibitors NaF [10 mM] and Na_3_VO_4_ [20 mM]) and quantified using a bicinchoninic acid (BCA) protein assay kit (#23225, Thermo Scientific). The samples were added with 6X loading dye and boiled at 97 °C for 5 min. An equal amount of 10-20 μg protein per line was used in the following SDS-PAGE and immunoblotting analysis. Anti-MAPK6 antibody was purchased from Abcam (ab53277). Antibodies against p-AKT T308 (#4056), p-AKT S473 (#4060), AKT (#2920), p-GSK3β S9 (#9558), RICTOR (#2114), HA-tag (#3724) and MK5 (#7419) were from Cell Signaling Technology (CST). Antibody against β-ACTIN (AC026) was purchased from ABclonal. Anti-FLAG (#200474) was from Agilent. Anti-MAPK4 antibody was purchased from ABGENT (#7298b).

### *In vitro* kinase assay

10x reaction buffer (500 mM Tris-HCl buffer [pH 7.5]; 1 mM EDTA; 2mM NaCl; 1% β-mercaptoethanol) and 5x Mg^2+^/ATP buffer (50 mM Magnesium Acetate, 2.5 mM ATP) were freshly prepared and stored at 4 °C for less than 24 hours. 1 × 10^6^ transient-transfected HEK293T cells or 3 × 10^6^ stable-infected PC3 cells were collected and inoculated in 1 ml of cell lysis buffer (20 mM Tris-HCl buffer [pH 7.5]; 150 mM NaCl; 2 mM EDTA; 1% Triton X-100; 1 mM NaF; 1 mM Na_3_VO_4_ and Proteinase inhibitor cocktail) at 4 °C for 30 min. The cell lysates were then centrifuged at 4°C at 13,800 g for 10 minutes. The supernatant was mixed with EZview Red ANTI-FLAG M2 Affinity Gel (Millipore Sigma) and rotated for 90 minutes at 4°C. After that, the ANTI-FLAG M2 Affinity Gel was washed with lysis buffer twice and with 1X reaction buffer three times. The reaction system contains 3 μl 10X reaction buffer, about 50 ng MAPK6 protein, 6 μl 5X Mg^2+^/ATP buffer, 100 ng recombinant His-AKT1 (inactive, Upstate/Millipore #14-279), at a final volume of 30 μl. Reaction conditions include incubation in a shaker (300 rpm) at 30 °C for 1h, tapping tube every 10 min, and finished with 37 °C for 20 min in a rotator.

### GST-pull down

The CDS of *MAPK6* was PCR amplified and cloned into the GTK3XF vector between EcoRI (5’) and SalI (3’) sites to generate the GST-MAPK6 fusion using the following primers: Forward: 5’- AAGGAATTCATGGCAGAGAAATTTGAAAGTCTCATG-3’; Reverse: 5’- AAGGTCGACTTAGTTCAGATGTTTCAGAATGCTGCTGTATG-3’. The GST-MAPK6 fusion protein was purified and subjected to pull-down assay as we previously reported (19). GST protein was used as control.

### Immunoprecipitation

Immunoprecipitation was performed using EZview Red ANTI-FLAG M2 Affinity Gel (Millipore Sigma), as we previously reported (19).

### Immunofluorescence

Cells were seeded on glass coverslip (VWR #89015-725) for 24-48h. Cells were then fixed with methanol for 15 min at 4 °C and washed with PBS three times. 0.3% Triton X-100 in PBS with 10% goat serum was used to increase the nuclear membrane permeability and block non-specific binding. The slides were then incubated with anti-AKT (CST #2920, 1:100) and anti-MAPK6 (Abcam ab53277, 1:100) antibodies overnight at 4°C. Alexa Fluor 488 goat anti-mouse IgG (1:500, Invitrogen #35502) and Alexa Fluor 594 goat anti-rabbit IgG (1:500, CST #8889) were used as the secondary antibody and incubated with cells at room temperature for 1 h. Finally, the slides were stained with DAPI and fixed on glass slides with mounting medium (#H-1000, Vector laboratories) and examined and photographed under an Olympus microscope.

### Xenograft

H1299 cells (1 × 10^6^) were s.c. injected into the lateral flanks of SCID mice (6-8 weeks old, purchased from Envigo). Mice began receiving 4 mg/ml Dox and 5% sucrose in drinking water on the day of tumor inoculation. Tumors were monitored every 2-3 days, and tumor volumes were calculated and recorded as 0.52 x length x width^2^. Tumors were harvested as indicated and weighed. Part of the tumor was used for protein extraction with RIPA buffer and subjected to western blots analysis.

### Analysis of human tumor molecular data sets

For pan-cancer survival analyses, we collected molecular data on 10,152 tumors of various histological subtypes (ACC project, n=79; BLCA, n=407; BRCA, n=1094; CESC, n=304; CHOL, n=36; CRC, n=619; DLBC, n=48; ESCA, n=184; GBM, n=159; HNSC, n=519; KICH, n=65; KIRC, n=533; KIRP, n=289; LAML, n=163; LGG, n=514; LIHC, n=370; LUAD, n=506; LUSC, n=495; MESO, n=86; OV, n=261; PAAD, n=178; PCPG, n=179; PRAD, n=497; SARC, n=259; SKCM, n=460; STAD, n=411; TGCT, n=134; THCA, n=503; THYM, n=119; UCEC, n=544; UCS, n=57; UVM, n=80) from The Cancer Genome Atlas (TCGA), for which both RNA-seq data (v2 platform) and patient survival data were available. Survival data were current as of March 31, 2016. Also, we analyzed a compendium of lung adenocarcinoma expression profiles (26) and a compendium of breast cancer expression profiles (27), compiled previously. Breast cancer survival data were capped at 15 years. The log-rank test evaluated the top 75% of MAPK6 expressing cases versus the other cases, while univariate Cox evaluated the log2 MAPK6 expression as a continuous variable.

### Statistics

Unpaired 2-tailed Student’s *t-*test was used to analyze the statistical significance in the cell-culture studies and paired 2-tailed Student’s *t*-test was used for comparing xenograft tumor weight. The proliferation assay and *in vivo* xenograft growth were analyzed using Two-way ANOVA. Two tailed tests and an α level of 0.05 were used for all statistical analyses. Survival analysis was carried out using log-rank test and univariate Cox (stratified log-rank test and stratified univariate Cox for the pan-cancer analysis, correcting for tumor type).

### Study approval

All animal studies were approved by the Institutional Animal Care and Use Committee of Baylor College of Medicine.

## Author Contributions

QC, WW, BD, TS, and FY designed the experiments, QC, BD, CJC, DDM, and FY wrote and revised the manuscript, QC, WW, and WZ performed the experiments, and QC, WW, BD, CJC, DDM, and FY analyzed the data.

## Acknowledgments

We thank Dr. Fengjun Chen for technical support. This research was supported by grants from the Department of Defense Congressionally Directed Medical Research Programs (W81XWH-17-1-0043 to F.Y.), the Cancer Prevention Research Institute of Texas (RP130651 to F.Y.), and the National Institutes of Health (CA125123 to C.J.C.). We also acknowledge the joint participation by Adrienne Helis Malvin Medical Research Foundation through its direct engagement in the continuous active conduct of medical research in conjunction with Baylor College of Medicine and the MAPK4 as a Novel Therapeutic Target for Human Cancers Cancer Program.

## Supplemental Information

**Figure S1.**
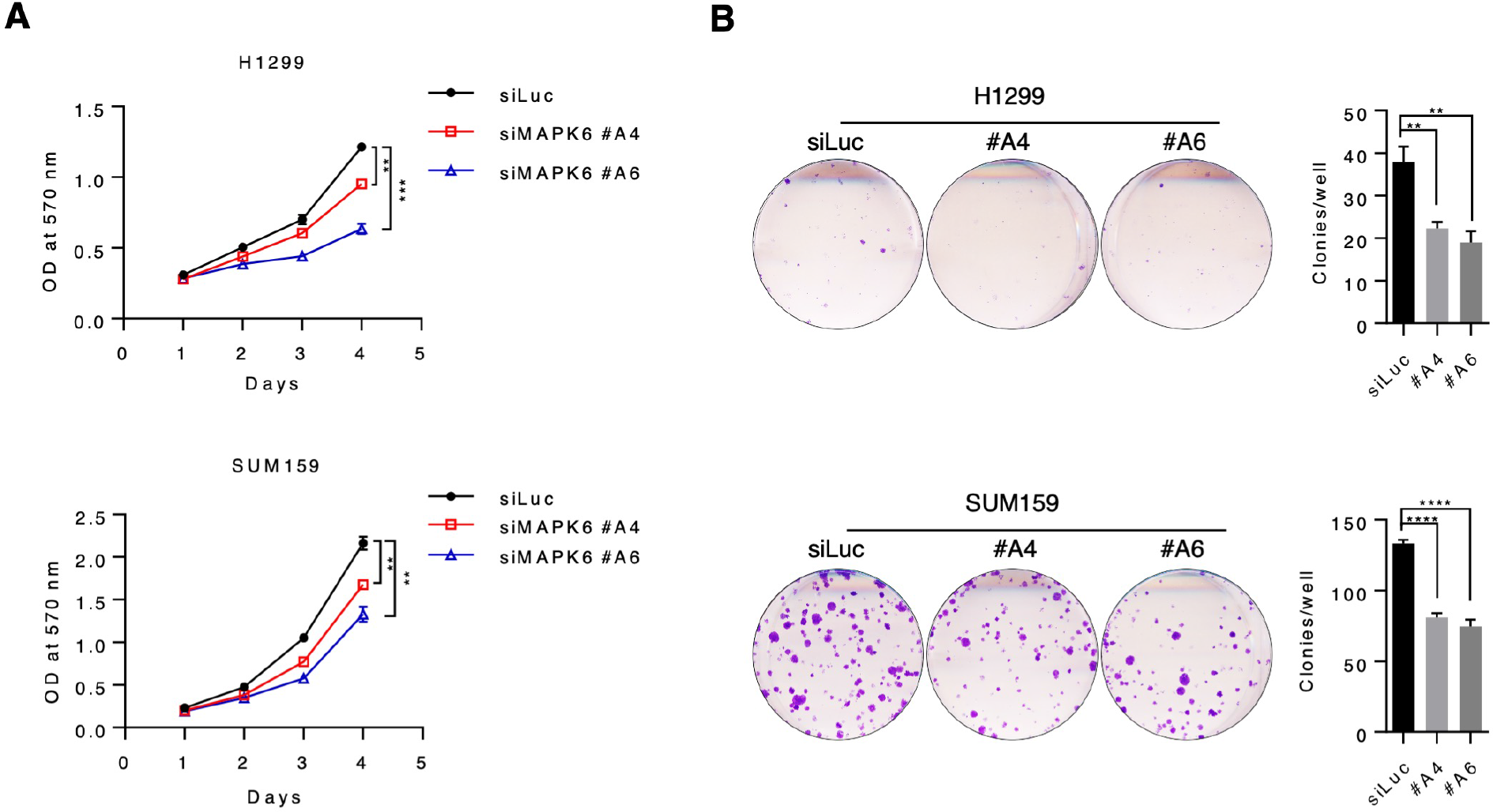
Knockdown of MAPK6 inhibits tumor cell growth. (**A**) The proliferation of the H1299 and SUM159 cells upon knockdown of MAPK6. (**B**) Plate colony formation of the H1299 and SUM159 cells upon knockdown of MAPK6 (mean ± SD, Student’s t-test). **: *P* < 0.01. ****: *P* < 0.0001. Data are representative of at least 3 independent experiments.

**Figure S2.**
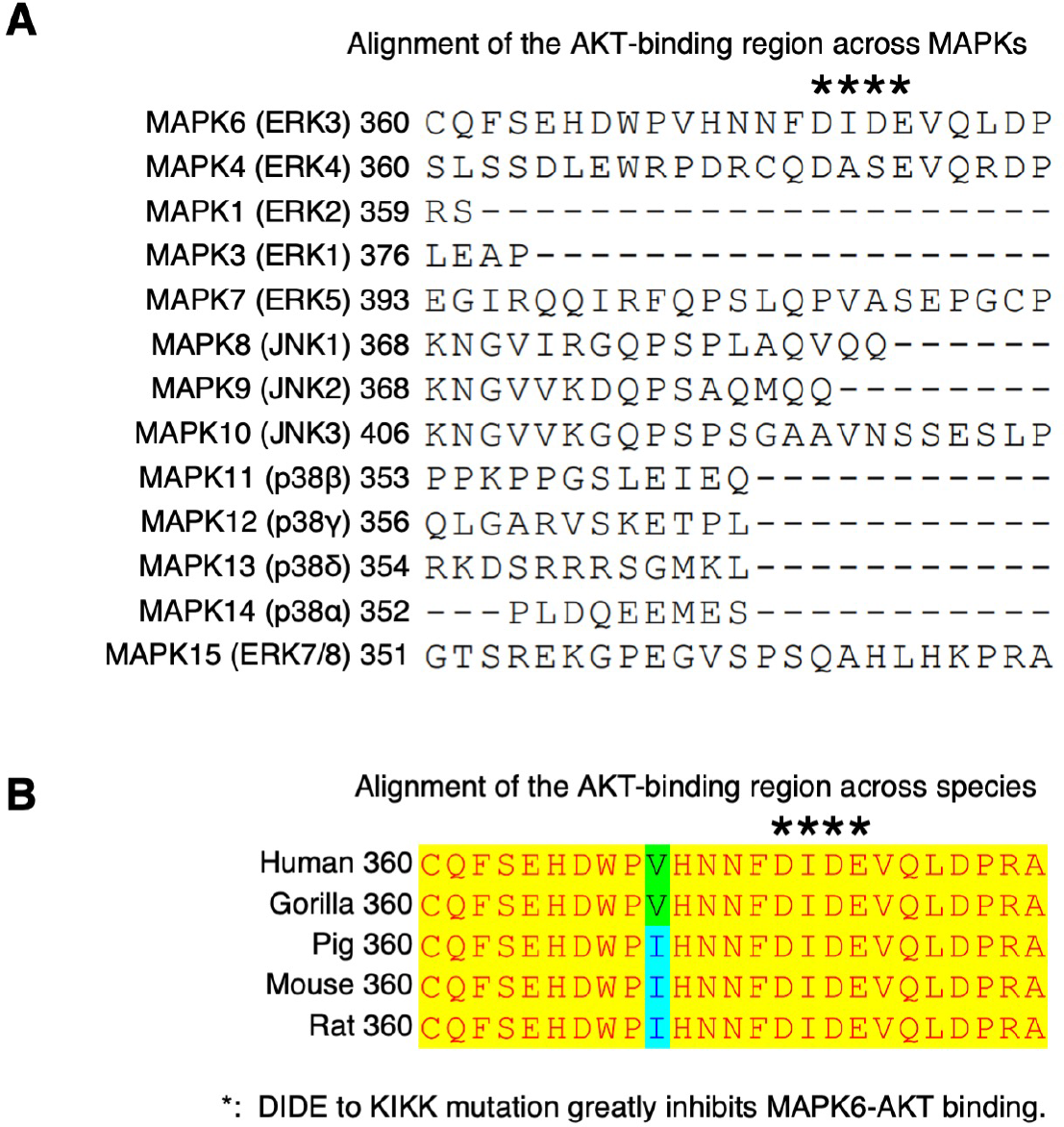
Alignments of MAPK6 protein sequences around the high affinity AKT-binding region across MAPKs and species. (**A**) There is little similarity across MAPKs. (**B**) the AKT-binding region is conserved across species.

